# Early Emergence of Cultural Differences in Audiovisual Speech Perception

**DOI:** 10.64898/2026.08.06.742174

**Authors:** Linlin Yan, Shuaike Hu, Yue Ding, Mengke Jin, Anna Krasotkina, Licai Ren, Shaoying Liu, Naiqi G. Xiao

## Abstract

Integrating auditory and visual cues is a hallmark of human speech perception, yet adults from East Asian backgrounds show less reliance on visual speech than their Western counterparts. The origins of this cultural difference, however, remain unknown. To investigate whether this divergence is established early in infancy, we examined audiovisual integration in 6– to 12– month-old White Canadian (n=111) and Chinese (n=115) infants using a novel paradigm measuring their perception of the McGurk effect. Across four experiments, we found a clear developmental divergence: Canadian infants showed a stable McGurk effect from 6 months onward, whereas Chinese infants showed a more protracted developmental trajectory, a cultural pattern that was further highlighted when their integration was challenged by other-race faces. These findings provide the first direct evidence that cultural differences in multisensory speech perception are established within the first year of life, suggesting that the brain’s strategy for binding sight and sound is shaped by early experience, with broad implications for theories of language acquisition and developmental science.

---

Human speech perception is inherently multimodal: listeners routinely integrate auditory input with visual cues from a speaker’s mouth movements. The McGurk effect illustrates this process. When a viewer sees a speaker articulating one syllable (e.g., /ga/) while hearing a different one (e.g., /ba/), they often perceive a third, fused syllable (e.g., /da/, McGurk & MacDonald, 1976). Beyond a compelling phenomenon, the McGurk effect indexes a core perceptual mechanism that underpins real-world communication. This mechanism is crucial for speech-in-noise comprehension (Ross et al., 2006; Sumby & Pollack, 1954), early language acquisition (Kuhl & Meltzoff, 1982; Lewkowicz & Hansen-Tift, 2012), and the development of robust phonological representations (Dodd, 1979). Therefore, understanding the factors that shape susceptibility to the illusion provides critical insights into the ontogeny of spoken language processing. While the phenomenon is easily engaged and widely recognized, susceptibility to the illusion is far from uniform. Research has reported stable individual differences in the McGurk effect that vary with age, language experience, and cultural background. For instance, adults from East Asian cultures (e.g., Japanese and Chinese) tend to show a weaker McGurk effect than Western adults, suggesting greater reliance on auditory cues (Sekiyama, 1997; Sekiyama & Tohkura, 1993, but see Magnotti et al., 2025). This cross-cultural variation raises a developmental question: when and how do culture-specific patterns of audiovisual integration emerge?

The investigation into these culture-specific patterns began with foundational research in adults. These studies revealed that individuals from East Asian cultures consistently exhibited a weaker McGurk effect than their Western counterparts (Sekiyama, 1997; Sekiyama & Tohkura, 1993). For instance, Japanese and Cantonese-speaking adults were found to be less influenced by incongruent visual information, more frequently reporting the veridical auditory syllable (Burnham & Lau, 1998; Sekiyama & Tohkura, 1993). While these findings established reliable cross-cultural differences in the adult perceptual outcome, it left the crucial question of its developmental origins entirely open.

A pivotal study by Sekiyama and Burnham (2008) provided the first evidence into the developmental trajectory of the cultural difference in the McGurk effect. They examined children aged 6 to 11 years and found that 6-year-old Japanese– and English-speaking children showed similarly weak susceptibility to the illusion. Between ages 6 and 8, however, English– speaking children began to rely more heavily on visual cues, a pattern that intensified into later childhood. The authors concluded that cultural differences in audiovisual integration become evident after 6 years of age. Yet their task required children to explicitly categorize fused sounds, which demands mature metalinguistic awareness and stable phonological categories. Because infants and preschoolers lack these abilities, the apparent absence of earlier divergence may reflect task demands rather than true developmental parity. The cultural split in audiovisual integration could well originate earlier, even during infancy.

Evidence from infant face-scanning pattern supports this possibility. By 7 months of age, British infants fixate more on the mouth of a talking face, whereas Japanese infants maintain a more central gaze (Geangu et al., 2016). This difference persists into toddlerhood (Haensel et al., 2020) and is consistent with attentional biases documented in Western and East Asian adults (Blais et al., 2008; Jack et al., 2009). This early gaze divergence has direct implications for audiovisual integration. Susceptibility to the McGurk effect is linked to the amount of visual attention directed at a speaker’s mouth (Gurler et al., 2015). If culturally specific face-scanning habits are already in place during infancy, the behavioural differences observed in later childhood may not emerge after age six but may instead follow from perceptual habits established in the first year of life.

Would the culture-related differences in the McGurk illusion already be established in preverbal infants? As briefly mentioned earlier, prior behavioral studies have provided converging evidence of McGurk-like audiovisual integration in Western infants (Burnham & Dodd, 1996, 2004; Desjardins & Werker, 2004; Rosenblum et al., 1997) and Asian infants (Ujiie et al., 2020, 2021). However, due to the inconsistency in experimental parameters, such as the syllables tested, it is challenging to draw direct comparisons between the findings from Eastern and Western infants. To the best of our knowledge, there’s no research specifically designed to directly investigate cultural variations in the McGurk effect during infancy. The cultural difference in infancy remains entirely unexplored.

To address this gap, we recruited 6– to 12-month-old infants from Mandarin-speaking families in China and from White monolingual English-speaking families in Canada. This age range was chosen because infants from both cultural backgrounds already show the McGurk effect before 6 months of age (Desjardins & Werker, 2004; Rosenblum et al., 1997; Ujiie et al., 2021), making the 6–12-month window suitable for detecting early cultural divergence. This period also coincides with perceptual narrowing, during which infants adapt to the specific properties of their native language environment (for a review, see Maurer & Werker, 2014). If culturally specific experience shapes audiovisual integration, its effects are likely to appear within this developmental window.

We designed a behavioural task in which infants hear a physically repeating syllable while viewing a face that alternates between articulating a McGurk-inducing syllable and remaining still. The task yields two diagnostically distinct looking-time patterns. First, if an infant consistently integrates the visual cues, the repeating sound should be perceived as an alternating sequence, because the articulating face alters the percept on every other syllable. Infants prefer alternating over repeating sequences (Gerken et al., 2011; Kidd et al., 2012), so consistent integration should produce longer looking times in McGurk trials than in non-McGurk control trials, where perception remains veridical and repetitive. Second, if integration is sporadic, the resulting percept is an unpredictable, random-like sequence. Infants prefer predictable repetition over such irregular patterns (Kidd et al., 2014), so inconsistent integration should produce shorter looking times in McGurk trials relative to controls. Comparing McGurk and non-McGurk looking times therefore distinguishes consistent integration, sporadic integration, and no integration without requiring explicit categorization.

Unlike traditional habituation methods, this approach minimizes memory demands and does not require assumptions about the specific illusory percept, thus offering a more direct measure of online audiovisual integration. We used this paradigm across three experiments. Experiments 1a and 1b validated the paradigm by confirming that infants’ looking times distinguished physically alternating from repeating sequences, and repeating from randomized sequences, respectively. Experiment 2 applied this paradigm to the central question, comparing audiovisual integration in White Canadian and Asian Chinese infants using familiar, own-race faces. Experiment 3 tested whether these cultural patterns hold when integration is challenged by unfamiliar, other-race faces (cf. Ujiie et al., 2021).

Several design choices controlled for potential confounds. All visual stimuli were generated using generative adversarial networks (GANs, Goodfellow et al., 2014) so that facial movements were identical across face identities and races (for independent evaluation of the naturalness of these faces, see Supplemental Materials). A caregiver questionnaire documented culturally specific caregiving arrangements (see Supplemental Materials). The Canadian and Chinese labs used identical eye-tracking systems, stimulus presentation setups, and recruitment protocols. This standardization ensures that any observed differences can be confidently attributed to cultural factors rather than procedural artifacts. Unless otherwise noted, Canadian infants were recruited from the [[CITY NAME]] region in Southern Ontario, Canada, whereas Chinese infants were recruited from the [[CITY NAME]] region on China’s eastern coast. The Chinese recruitment region is racially homogeneous (>99.99% Han Chinese). The same recruitment procedures and inclusion criteria were applied across Experiments 1–3.

## Experiment 1a

Before investigating audiovisual integration, we first conducted two benchmark experiments to validate our core methodological premise that infants’ looking times in this paradigm would reliably reflect their perception of different auditory sequences. The central hypothesis of our main experiments is that consistent integration of a McGurk stimulus would create a perceptually alternating sound sequence, which infants would prefer over a repeating one. Experiment 1a was therefore designed to confirm this baseline preference by measuring looking times to a physically alternating sequence versus a repeating one.

Because inconsistent audiovisual integration would produce an unpredictable percept rather than a stable alternating sequence, Experiment 1b established infants’ response to randomized auditory sequences. Together, Experiments 1a and 1b provide benchmarks for interpreting consistent, sporadic, and absent audiovisual integration in subsequent experiments. Importantly, each benchmark experiment was conducted with both White Canadian and Chinese infants to validate the procedure across laboratory contexts. These experiments served a methodological purpose: to confirm comparable baseline sensitivity to auditory sequence structure before testing cross-cultural differences in audiovisual integration.

### Participants

Two cohorts of infants participated in Experiment 1a. The first cohort consisted of 23 full-term White Canadian infants (11 females) with normal vision and hearing. Their ages ranged from 183 to 379 days old (*M* = 9.79 months, *SD* = 1.87 months). All infants came from monolingual English-speaking families and had minimal exposure to East-Asian faces in daily life according to parental reports. The second cohort consisted of 20 full-term Chinese Asian infants (8 females) with normal vision and hearing. Their ages ranged from 151 to 412 days (*M* = 9.43 months, *SD* = 2.97 months).

All experimental protocols (Experiments 1, 2, & 3), including the procedures for obtaining informed consent, were reviewed and approved by the Research Ethics Board of the host institute in Canada and the Institutional Ethics Board of the host institute in China. Caregivers provided written informed consent, and participants received a gift in exchange for their participation.

### Stimuli

We created the experimental stimuli by first video recording a female actress articulating four syllables (/ba/, /pa/, /ga/, and /ka/) with a neutral expression. Each articulation was trimmed to a 1-second clip. The audio component of these clips was processed to remove background noise and normalize loudness. For the visual component, we used generative adversarial networks (GANs; Goodfellow et al., 2014) architecture to transfer an actress’s facial movements onto 16 photorealistic faces (8 White, 8 East-Asian). Specifically, we utilized the DeepFaceLab 2.0 framework (Perov et al., 2020), augmented with custom code to control the temporal synchronization and visual fidelity. This process minimized potential bias from variations in facial movements by ensuring identical, artifact-free facial motions across all faces. To address concerns regarding ecological validity, an adult validation study (detailed in Supplemental Materials) confirmed that these synthetic stimuli were perceived as highly natural, with ratings statistically indistinguishable from real human videos. We then standardized the visual stimuli by removing the hair, normalizing the face size (16.3 × 16.3 cm on the screen), and placing them against a uniform light gray background, while preserving other facial features to maintain a natural appearance.

### Procedure

To measure infants’ auditory perception via looking time, we created two sound sequences: ***alternating***, where two distinct sounds were presented in alternation, and **repeating**, where a single sound was presented throughout the trial. The specific sounds used in each trial were randomly selected from the four syllables. Each sequence consisted of 20 one-second sounds.

In each trial, the auditory stimuli were paired with visual presentations of a face. For the 1st, 3rd, and all subsequent odd-numbered sounds, a video of an articulating face was shown, with the mouth movements synchronized to the sound. For the 2nd, 4th, and all subsequent even– numbered sounds, a still image of the face (the first frame of the video) was displayed. This created a “moving-still” visual pattern synchronized with the auditory sequence, resulting in a trial where infants observed an alternating pattern of articulating and static faces while hearing either an alternating or repeating sound sequence. The order of the alternating and repeating trials was randomized across participants.

To assess infants’ auditory perception, we measured their on-screen looking time using an EyeLink 1000 Plus eye-tracker (500 Hz sampling rate; SR Research, Canada). Infants sat on their caregivers’ laps in a sound-attenuated testing room throughout the experiment. Parents were instructed not to interfere with their infants’ looking behaviors prior to the experiment. Infants sat approximately 60 to 80 cm away from the 55 × 31 cm^2^ (25-inches) computer monitor, where we used Psychtoolbox (3.19.8) software to play the visual stimuli. The experiment started with an infant-controlled calibration program to ensure eye tracking precision and accuracy. During calibration, a cartoon figure was presented sequentially at five locations (four corners and the center) with a rewarding rattling sound. Calibration was targeted for an average spatial error below 1° of visual angle. Our validation records confirmed data quality across both laboratory sites: the mean validation error across studies for all Canadian infants was 0.93° (*SD* = 0.69°), and for the Chinese infants, it was 0.76° (*SD* = 0.28°). At the beginning of each trial, an attention-getter (i.e., a colorful bouncing ball) located at the center of the monitor directed infants’ attention back to the screen. Experiments 1a and 1b each consisted of a maximum of 16 trials.

### Data Analysis

Raw eye-tracking data were processed into fixation data using EyeLink DataViewer software (Version 4.4.1). Trials were excluded if total fixation time was below 1000 ms, corresponding to less than one complete audiovisual event. For each participant, mean looking time was then calculated for each trial type and used as the primary dependent measure. The same preprocessing and exclusion criteria were applied across Experiments 1, 2, and 3.

Paired-sample and independent-samples t-tests were used to compare looking times within and between groups, respectively. All tests were two-tailed with *α* = .05. Bayes factors (*BF*₁₀) were computed for all t-tests using the BayesFactor package in R (Version 4.6.1; Morey & Rouder, 2026) to quantify evidence for the alternative hypothesis over the null. Following Jeffreys’ (1961) conventions, *BF*₁₀ values of 1–3 were interpreted as anecdotal evidence, 3–10 as moderate evidence, 10–30 as strong evidence, 30–100 as very strong evidence, and >100 as decisive evidence for the alternative hypothesis.

### Results and Discussion

As shown in Figure 1, Canadian infants exhibited significantly longer looking times during alternating trials (*M* = 14.16, *SD* = 3.29) compared to repeating trials (*M* = 13.35, *SD* = 3.21; paired-sample *t*(22) = 2.18, *p* = .04, Cohen’s *d* = 0.45, *BF*_10_ = 1.57). Similarly, Chinese infants showed significantly longer looking times during alternating trials (*M* = 15.03, *SD* = 3.09) relative to repeating trials (*M* = 14.00, *SD* = 3.44; paired-sample *t*(19) = 2.89, *p* = .009, Cohen’s *d* = 0.65, *BF*_10_ = 5.40). Together, these results confirm that infants discriminated the benchmark auditory sequences, validating the paradigm for subsequent audiovisual integration experiments. Nevertheless, the overall pattern confirms the validity of looking time as a robust measure of auditory discrimination in this paradigm.

**Figure 1.**
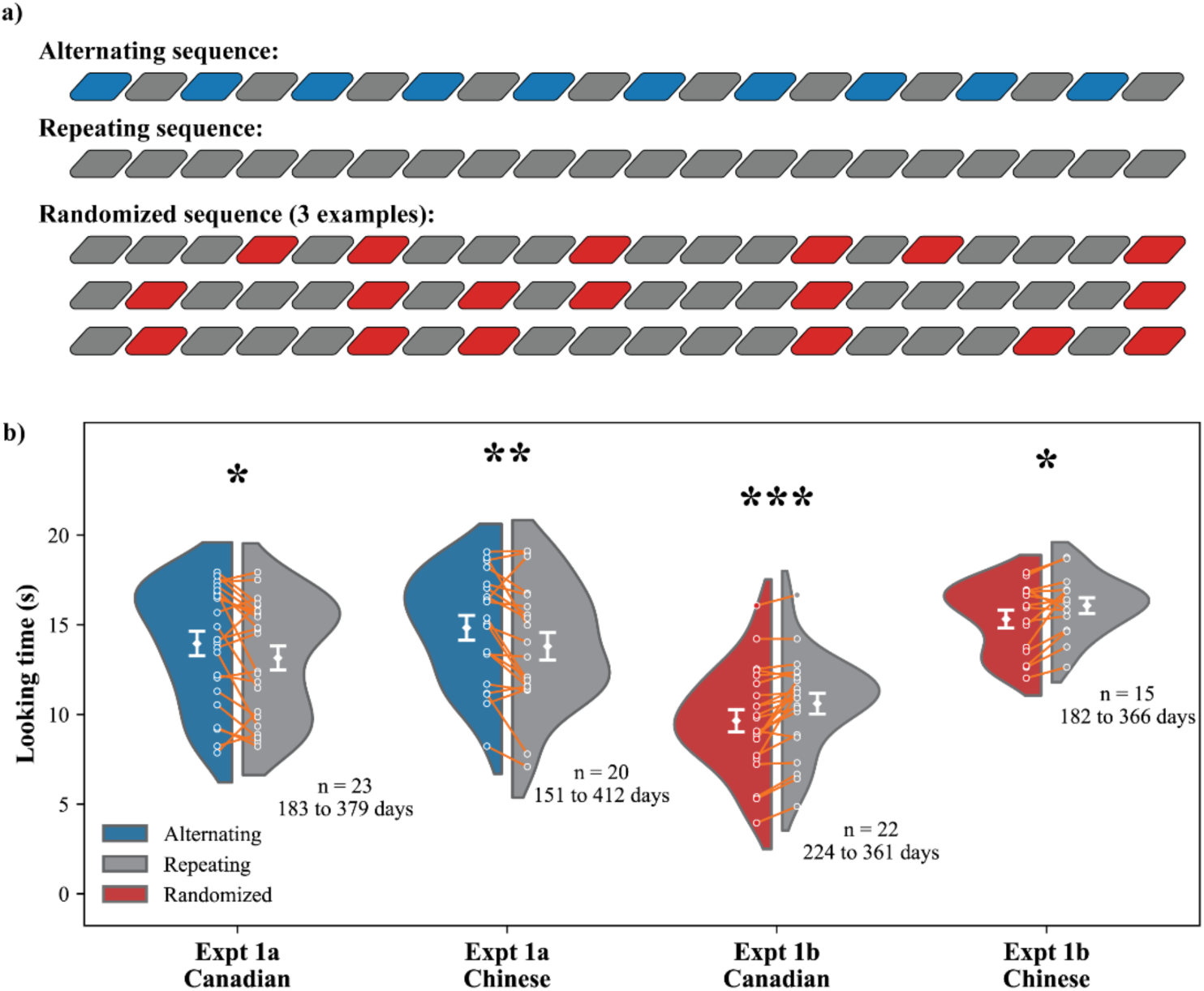
Mean looking time for different trial types of the in Experiments 1a and 1b. The white diamond shapes represent the mean looking time. Error bars represent one standard error from the mean. Each participant’s looking time was annotated with tiny white dots, and the thin orange lines connecting the dots indicated the looking time of the same participant. The kernel distributions (the split violin shapes) represent estimated distribution of the mean looking time across participants within each condition and cohort. The asterisks indicate statistically significant difference in looking time between conditions.

Looking-time difference scores (alternating − repeating) did not differ between cultures, *t*(41) = −0.44, *p* = .66, suggesting that the preference for alternating sequences reflects domain– general perceptual mechanisms shared across cultures.

## Experiment 1b

### Participants

Two cohorts of infants participated in Experiment 1b. The first cohort consisted of 22 full-term White Canadian infants (12 females) with normal vision and hearing, ranging in age from 224 to 361 days (*M* = 9.62 months, *SD* = 1.62 months). The second cohort comprised 15 full-term Chinese infants (9 females) with normal vision and hearing. Their ages ranged from 182 to 366 days (*M* = 8.52 months, *SD* = 2.07 months). These infants were recruited from the same regions along China’s eastern coast as in Experiment 1a.

### Stimuli & Procedure

In Experiment 1b, we introduced randomly ordered sound sequences to contrast with the repeating pattern. These were created by modifying the alternating sequences from Experiment 1a: the sound at each even-numbered position (2nd, 4th, etc.), which co-occurred with a still face, was replaced by the sound from the preceding odd-numbered position (1st, 3rd, etc.). This ensured that the presented sound always matched the syllable articulated by the moving face in the preceding visual event, creating a less predictable auditory stream.

From a larger pool of generated sequences, we selected those with the Language of Thought (LoT) complexity values (Al Roumi et al., 2023) for the experiment. These random– sequence trials were presented with repetition trials, which were identical to those used in Experiment 1a. The order of the randomized and repeating trials was randomized across participants.

### Results and Discussion

For the White Canadian cohort, infants looked significantly longer during repetition trials (*M* = 10.81, *SD* = 2.68) than during randomized trials (*M* = 9.85, *SD* = 2.92; paired-sample *t*(21) = 4.17, *p* < .001, Cohen’s *d* = 0.89, *BF*_10_ = 75.16; Figure 1). Similarly, the Chinese cohort showed significantly longer looking times during repetition trials (*M* = 16.27, *SD* = 1.69) compared to randomized trials (*M* = 15.52, *SD* = 1.95; paired-sample *t*(15) = 2.77, *p* = .015, Cohen’s *d* = 0.71, *BF*_10_ = 3.99; Figure 1). Take together, these findings indicate a consistent perceptual preference for predictable, repeating auditory sequences over random presentations in both groups.

An independent-samples *t*-test comparing the looking-time differences scores between the two cohorts revealed no significant cross-cultural difference, *t*(35) = –0.57, *p* = .57, confirming that the overall magnitude of this preference did not vary between populations.

In summary, Experiments 1a and 1b showed similar looking preferences across Canadian and Chinese infants, validating looking time as a measure of auditory perception. The absence of cultural differences suggests that these preferences reflect domain-general statistical learning mechanisms (Al Roumi et al., 2023; Kidd et al., 2012, 2014), providing a baseline for interpreting audiovisual integration in the subsequent experiments.

## Experiment 2

### Sample size estimation

The sample size of Experiment 2 was determined with consideration of both statistical power and the practical constraints typical of infant eye-tracking research. Initial pilot data (n = 4, aged 279 to 365 days) from an experiment with identical methodology yielded a large effect size (Cohen’s *d* = 1.58). A power analysis based on this estimate (α = .05, power = .90, two– tailed paired-sample *t*-test) using the R package “pwr” indicated that approximately 6 participants would be sufficient to detect a reliable McGurk effect in infants. Similarly, a second power analysis based on pilot data from 10 participants (aged 185–365 days) showing a correlation of *r* = .86 between age and the McGurk effect (α = .05, power = .95, two-tailed) suggested that approximately 10 participants would suffice to detect age-related changes.

However, we acknowledge that these pilot-based estimates are likely unstable given the relative small sample sizes (*n* = 4 and *n* = 10), and therefore cannot serve as a reliable basis for sample size determination. It should be noted that demonstrating this within-subject effect was merely a prerequisite. The primary theoretical goal of Experiment 2 was to test between-subject cultural differences and Culture × Age Group interactions. Because such comparisons involve greater between-subject variability than within-subject condition contrasts, and because infant eye-tracking studies typically involve high variability and attrition, we adopted a conservative recruitment strategy. We therefore targeted approximately 45 infants per cultural group, a sample size that is consistent with, and in many cases exceeds, previous infant audiovisual integration studies (e.g., Kushnerenko et al., 2013; Burnham & Dodd, 2004). This larger sample was intended to provide sufficient robustness for the primary cross-cultural and age-split analyses rather than to rely on unstable pilot-derived effect-size estimates.

### Participants

Forty-five full-term English-learning Canadian infants (10 females, 38 White and 7 mixed-race infants with one White primary caregiver) with normal vision and hearing participated in the current experiment after caregivers provided informed consent. All infants were from monolingual families. The participants were from 183 to 368 days old (*M* = 8.40 months, *SD* = 1.54 months). According to parental reports, all participating infants had minimal exposure to East-Asian faces in daily life. To ensure that the White face stimuli represented infants’ predominant everyday face experience, we recruited White Canadian infants whose primary caregiving environment was predominantly White (i.e., both parents or the primary caregivers were White). Nineteen additional infants participated but were excluded from the data analysis because of failure to complete the experimental procedure due to fussiness (n = 11), calibration failure (n = 5), parental interference (n = 1), equipment glitch (n = 1), caregivers’ race being non-White (n = 2), or performance that was more than two standard deviations from the group mean (n = 3). All participating families were from urban and suburban regions.

Forty-seven full-term Mandarin-learning Chinese infants (24 females) with normal vision and hearing participated after caregivers gave informed consent. The participants were from 182 to 375 days old (*M* = 9.04 months, *SD* = 2.23 months). Additional eighteen infants’ data were excluded from data analyses because of fussiness (n = 8), calibration failure (n = 6), equipment glitch (n = 1), parental interference (n = 2), and performance that was more than two standard deviations from the mean (n = 1). All participating families were from urban and suburban regions.

### Stimuli

To probe infants’ audiovisual integration, Experiment 2 employed a similar paradigm but introduced two distinct types of stimuli: **McGurk** and **non-McGurk**. The **McGurk** stimuli featured audiovisual pairings known to elicit an illusory auditory percept (e.g., McGurk & MacDonald, 1976; Rosenblum et al., 1997). For example, the visual syllable /ga/ was paired with the audio syllable /ba/, which typically results in the perception of /da/. The **non-McGurk** stimuli served as a control and were created by reversing the components of the McGurk pairs (e.g., visual /ba/ with audio /ga/, Kushnerenko et al., 2008). This combination is unlikely to produce an illusion, with perception typically remaining veridical to the audio input.

This counterbalanced design offers two key advantages. First, since both conditions involve a mismatch between the auditory and visual streams, any differential response can be attributed to the specific nature of the audiovisual integration rather than to sensory incongruity alone. Second, by presenting the exact same set of audio and visual syllables across both trial types, the design controls for potential infant preferences for any particular syllable.

In Experiment 2, all participants were presented with familiar-race, female faces. Specifically, White Canadian infants viewed White female faces, and Chinese infants viewed East Asian female faces. This established a baseline measure of audiovisual integration where visual speech cues are provided by a familiar face type.

We used the McGurk and non-McGurk pairs to create the McGurk and non-McGurk trials, respectively. Video demonstrations of the study stimuli are available on the Open Science Framework (OSF) website at the following link: https://osf.io/a6qwf/?view_only=d678127560da45a198f69260a84abd11.

**Table 1.**
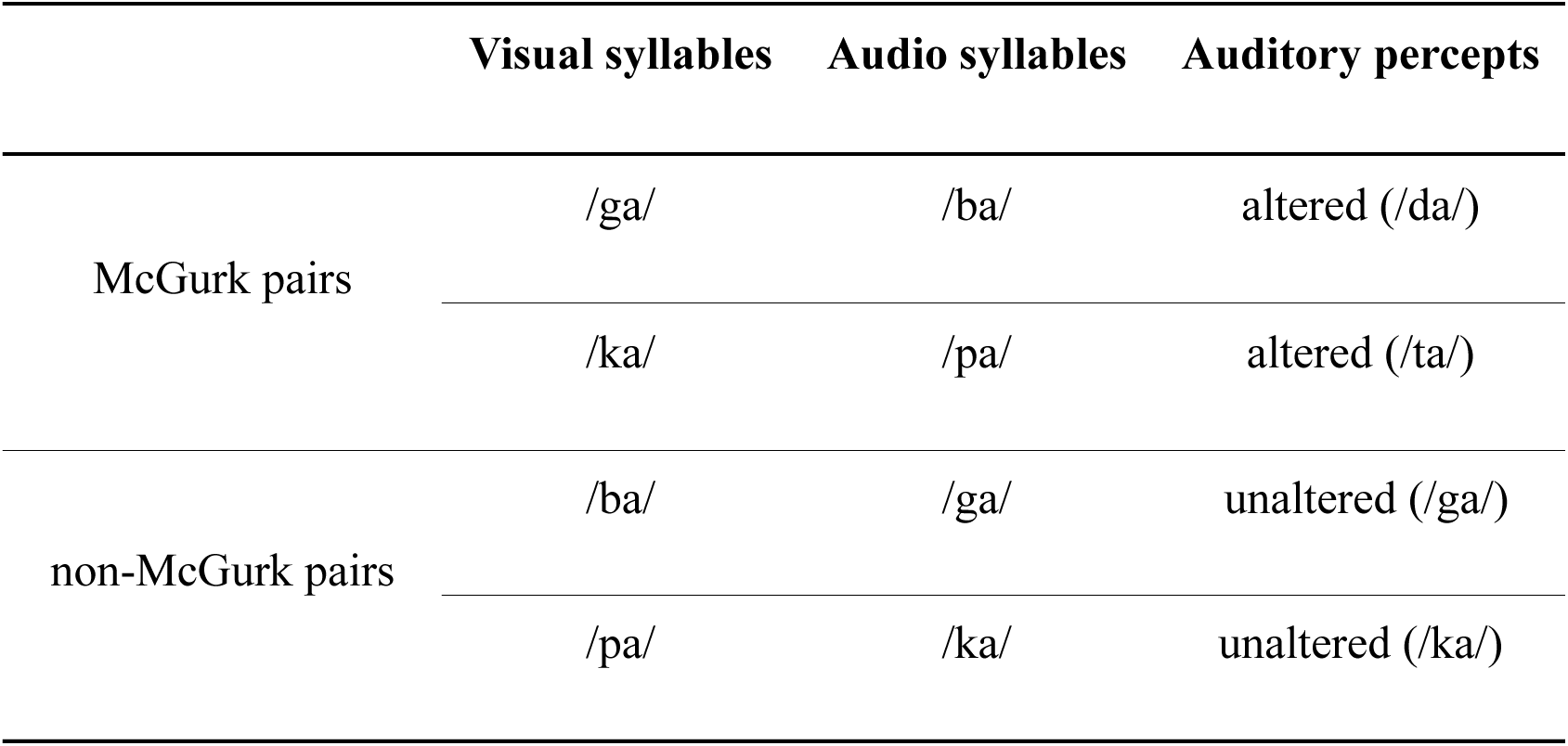
The audio and visual combinations of the McGurk and non-McGurk pairs used in the current study.

|  | Visual syllables | Audio syllables | Auditory percepts |
| --- | --- | --- | --- |
| McGurk pairs | /ga/ | /ba/ | altered (/da/) |
|  | /ka/ | /pa/ | altered (/ta/) |
| non-McGurk pairs | /ba/ | /ga/ | unaltered (/ga/) |
|  | /pa/ | /ka/ | unaltered (/ka/) |

### Procedure

Infants were presented with up to 16 trials, composed of **McGurk** and **non-McGurk trial** types, to assess their audiovisual integration. In each **McGurk trial**, a single McGurk audiovisual pair was presented 20 times. The visual presentation followed the “moving-still” paradigm established in Experiment 1: the articulating face was shown on odd-numbered iterations (1st, 3rd, etc.), while a static image of the face was shown on even-numbered iterations (2nd, 4th, etc.). The critical manipulation is that if infants integrate the visual information, an illusory auditory percept should arise only when the face is moving. This would transform a physically repeating audio stream into a perceived alternating sound sequence.

The **non-McGurk trials** were structurally identical but utilized the non-McGurk pairs. Since these pairings are unlikely to alter auditory perception, infants were expected to perceive the physically repeating audio stream as a simple repeating sound sequence throughout the trial. Figure 2a provides a schematic of the stimuli and their expected perceptual outcomes for both trial types.

**Figure 2.**
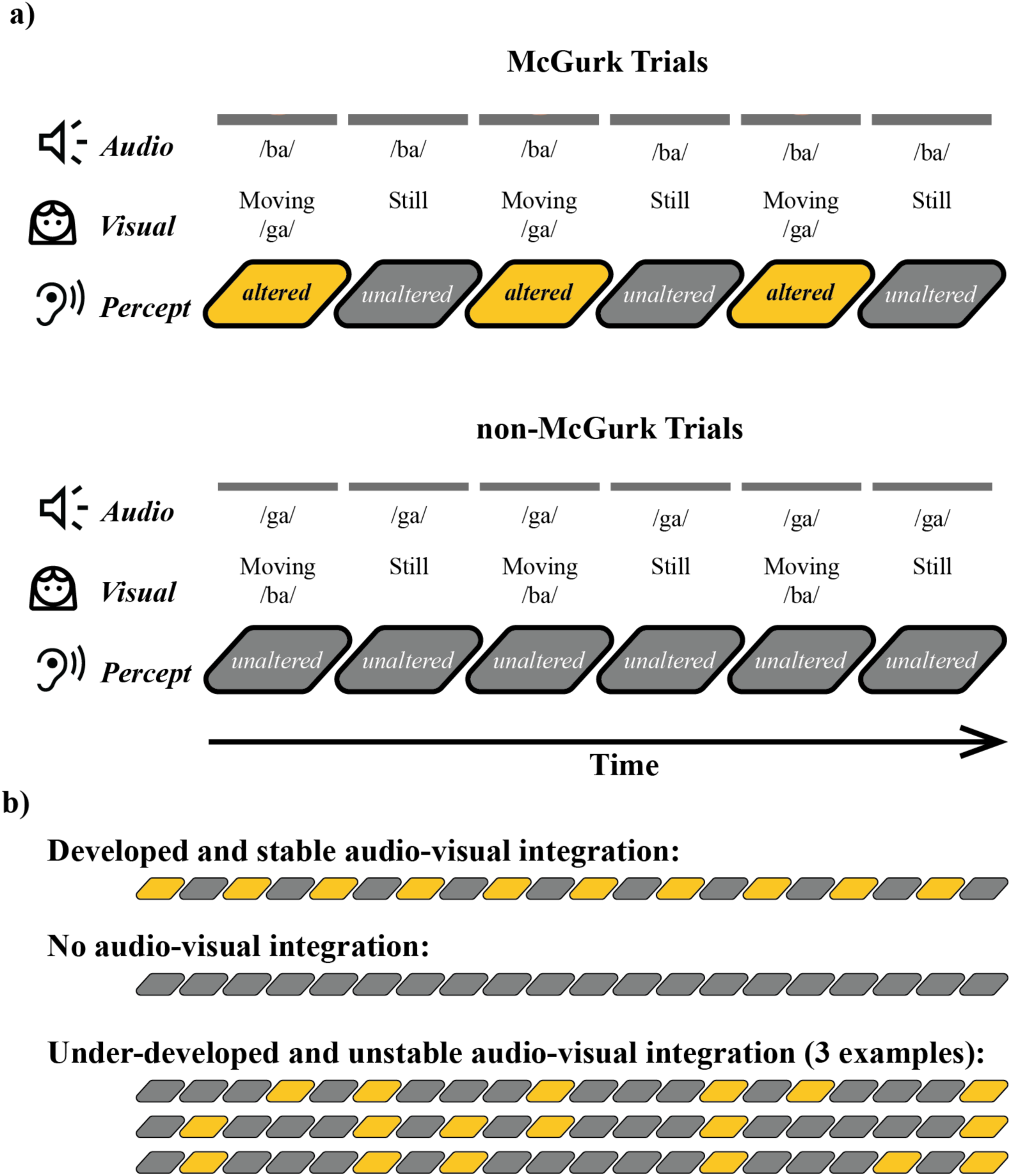
a) Schematic illustration of the experimental procedure in the McGurk and non– McGurk trials. We used one White face among the eight White faces and two audiovisual stimuli pairs (i.e., visual /ga/-audio /ba/ and visual /ba/-audio /ga/) for demonstration purposes. b) Three possible auditory perceptual outcomes of the McGurk trials. The yellow boxes represent altered auditory perception, and the grey boxes represent unaltered auditory perception.

This design allows us to interpret infants’ looking time patterns based on the consistency of their audiovisual integration. Drawing from research showing that the McGurk effect is not an all-or-nothing phenomenon (e.g., Sekiyama & Burnham, 2008), we conceptualize infant integration as a continuous ability. This leads to three distinct, testable hypotheses regarding their looking behavior (Figure 2b):

1. **Consistent Integration:** If infants reliably integrate visual articulatory information, they should perceive the McGurk trials as a structured, alternating sequence of two sounds, while perceiving the non-McGurk trials as a simple repetition of one sound. Given that infants show a preference for structured, complex sequences over simple, repetitive ones (Al Roumi et al., 2023; Kidd et al., 2012, 2014), we predicted that consistent integration would result in **longer looking times in McGurk than in non-McGurk trials**.
2. **Sporadic Integration:** If infants’ audiovisual integration is still developing, the integration of visual speech might occur inconsistently. This would cause the McGurk trials to be perceived as an irregular, unpredictable mixture of altered and unaltered sounds. Because infants demonstrate reduced interest in random or unpredictable sequences that lack a learnable pattern (Kidd et al., 2012, 2014), we predicted that this partial integration would lead to **shorter looking times in McGurk than in non– McGurk trials**.
3. **No Integration:** If infants are unable to integrate the visual articulatory information at all, their auditory perception will be driven solely by the physical sound input. In this scenario, they would perceive a repeating sound sequence in both the McGurk and non– McGurk trials, leading to no significant difference in looking times between the two conditions. While such a null finding aligns with a lack of robust audiovisual integration, it could also stem from alternative factors such as insufficient statistical power or measurement noise.

To ensure that looking time reflected auditory perception, we used a gaze-contingent design in which the face stimulus played only while infants fixated within the face region. Looking away for more than 300 ms paused the video until gaze returned, ensuring that looking time reflected active engagement (Xiao et al., 2023).

The study included four blocks. There were four trials within each block: two McGurk trials (visual /ga/-audio /ba/ and visual /ka/-audio /pa/) and two non-McGurk trials (visual /ba/– audio /ga/ and visual /pa/-audio /ka/). The trials within each block were played to the infants in a randomized order, and no identical trial was presented twice in a row. The experiment ended after infants completed four blocks of four trials (16 trials total), or earlier between trials if the infant became excessively fussy, if parents requested termination, or if the experimenter judged that continuation was no longer appropriate for the infant’s comfort. Once a trial began, the full 20-second audiovisual sequence was always presented in its entirety; the program did not allow within-trial termination. Manual session termination required the experimenter to press and hold the delete key until the onset of the next trial, ensuring that termination could occur only between trials and could not be triggered accidentally during stimulus presentation. There were 8 distinct face identities. These identities were randomly arranged to fill all 16 trials depicting the four types of articulatory movements. The presentation order of all identities and trial types was fully randomized.

### Results and Discussion

On average, Canadian infants completed 14.6 trials (*SD* = 2.92), and Chinese infants completed 9.62 trials (*SD* = 2.35). Preliminary analyses revealed that infants’ looking times did not significantly differ between the two syllable pairings (/BaGa/ and /PaKa/) within each condition. Therefore, for the primary analysis, looking time data were collapsed across syllable pairings within their respective McGurk and non-McGurk trial types. To examine whether differences in trial completion or overall looking duration accounted for the cultural– developmental pattern, we conducted additional trial-level mixed-effects and exposure-matched analyses (see Supplemental Materials). The results showed that the observed effects below were not attributable to differences in trial dynamics.

### Audiovisual integration in White Canadian infants

To assess audiovisual integration in White Canadian infants, a paired-sample *t*-test was conducted on the looking times for McGurk versus non-McGurk trials. As shown in Figure 3a, infants looked significantly longer during McGurk trials (*M* = 11.68, *SD* = 2.71) than non– McGurk trials (*M* = 10.95, *SD* = 3.32), paired-sample *t*(44) = 3.52, *p* = .001, Cohen’s *d* = 0.53, *BF*_10_ = 29.19. This result aligns with the consistent integration hypothesis, indicating that infants perceived the physically repeating audio in the McGurk trials as an alternating sequence, thereby demonstrating that articulatory facial movements altered their perception of the audio syllables.

**Figure 3.**
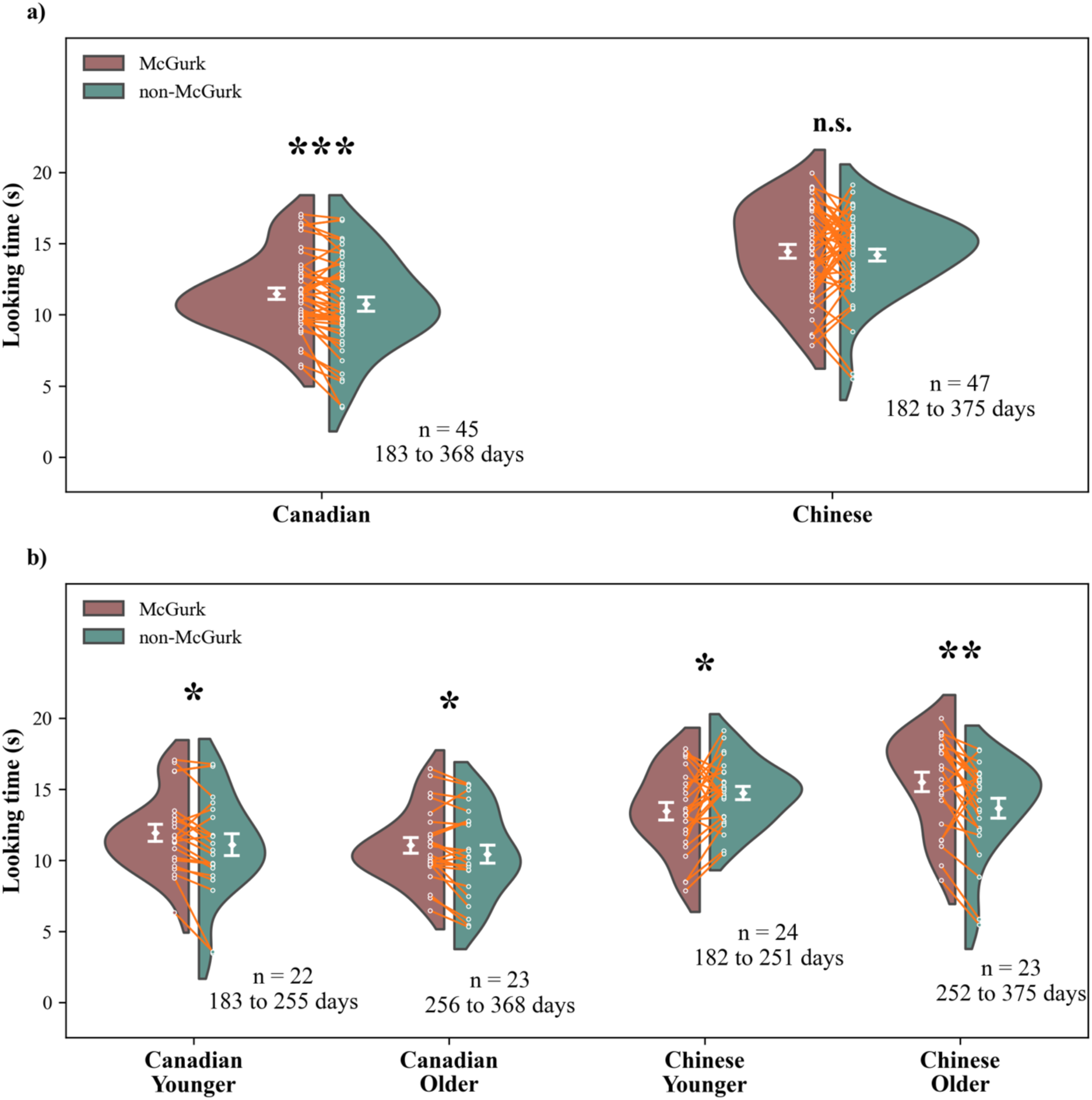
Mean looking time for Canadian and Chinese infants in McGurk and non-McGurk trials (Experiment 2). (a) Results are shown for the full sample in each cultural group. (b) Results are shown for infants separated into younger and older age cohorts. The white diamond shapes represent the mean looking time. Error bars represent one standard error from the mean. Each participant’s looking time was annotated with tiny white dots, and the thin orange lines connecting the dots indicated the looking time of the same participant. The kernel distributions (the split violent shapes) represent estimated distribution of the mean looking time across participants within each condition and cohort. The asterisks indicate statistically significant difference in looking time between the McGurk and non-McGurk trials.

To examine the developmental trajectory of this ability, we performed a Pearson’s correlation between infant age (in days) and a looking-time difference score (McGurk minus non-McGurk). The analysis revealed no significant age-related change in performance, *r*(43) = –.14, *p* = .349, *BF*_10_ = 0.49. This suggests that for Canadian infants, audiovisual integration is robustly established and stable across the 6– to 12-month age range. Furthermore, this stability implies that the ability likely emerges prior to 6 months of age, a finding consistent with previous research (Burnham & Dodd, 2004; Rosenblum et al., 1997).

### Age-related change in audiovisual integration in Chinese infants

An initial paired-sample *t*-test comparing McGurk and non-McGurk trials for the entire Chinese cohort revealed no significant overall difference in looking time (*M*_McGurk_ = 14.65, *SD* = 3.26; *M*_non-McGurk_ = 14.40, *SD* = 2.90; *t*(46) = 0.55, *p* = .585, *Cohen’s d* = 0.08, *BF*_10_ = 0.18).

We hypothesized that this null finding, derived from a sample spanning a wide age range (182 to 375 days), masked significant developmental changes. Specifically, based on our predictions, we expected younger infants with sporadic integration to look longer at non-McGurk trials, whereas older infants with consistent integration would look longer at McGurk trials; averaging these opposing patterns would yield a non-significant result. To test this hypothesis, we examined the age-related change in the looking-time difference score (McGurk minus non– McGurk). A Pearson correlation confirmed a significant positive age-related increase in preference for the McGurk trials, *r*(45) = .32, *p* = .026, *BF*_10_ = 3.03.

To further delineate this developmental change, infants were split into younger (182 to 252 days) and older (253 to 375 days) cohorts based on the median age (252 days). For the younger cohort (*n* = 24), infants looked significantly longer at non-McGurk trials (*M* = 14.92, *SD* = 2.31) compared to McGurk trials (*M* = 13.64, *SD* = 2.94), *t*(23) = 2.26, *p* = .034, Cohen’s *d* = 0.46, *BF*_10_ = 1.77. This finding supports the sporadic integration hypothesis. In contrast, the older cohort (n = 23) showed the opposite pattern, looking significantly longer at McGurk trials (*M* = 15.70, *SD* = 3.32) than non-McGurk trials (*M* = 13.85, *SD* = 3.38), *t*(22) = 3.33, *p* < .01, Cohen’s *d* = 0.69, *BF*_10_ = 13.48 (Figure 3b). This preference for the McGurk condition aligns with the consistent integration hypothesis. Collectively, these results demonstrate that the capacity for audiovisual integration undergoes a substantial and directional developmental change between 6 and 12 months of age in Chinese infants, shifting from a less consistent integration pattern to a robust one.

### Cultural difference in the development of audiovisual integration

To directly compare the developmental trajectories between cultures, we first confirmed the performance pattern for the Canadian infants. Using a median age split (256 days), we found that both younger (183 to 255 days) and older (256 to 368 days) Canadian infants looked significantly longer at McGurk (younger: *M* = 12.13, *SD* = 2.80; older: *M* = 11.28, *SD* = 2.61) than at non-McGurk trials (younger: *M* = 11.25, *SD* = 3.58; older: *M* = 10.63, *SD* = 3.10; younger: paired-sample *t*(21) = 2.44, *p* = .024, Cohen’s *d* = 0.52, *BF*_10_ = 2.43; older: paired– sample *t*(22) = 2.59, *p* = .017, Cohen’s *d* = 0.54, *BF*_10_ = 3.21). With this stable pattern of robust integration established for the Canadian cohort, we then conducted a 2 (age group: younger, older) × 2 (country: Canada, China) Analysis of Variance (ANOVA) on the looking-time difference score (McGurk minus non-McGurk) to formally test for a cultural difference in the developmental trajectory.

The ANOVA revealed a significant interaction between age group and country, *F*(1, 88) = 13.56, *p* < .001, *partial Eta Squared* = .13, *BF*_10_ = 67.09, indicating a cultural difference in developmental trajectories. Post-hoc comparisons using Tukey’s HSD test were conducted to deconstruct this interaction. These tests revealed that the younger Chinese infants differed significantly from their Canadian counterparts (*p* = .007), whereas this cultural difference was not present in the older cohorts (*p* = .233). Furthermore, the analysis confirmed a significant developmental shift within the Chinese cohort between the younger and older groups (*p* < .001), a pattern that was absent for the Canadian cohort, whose performance remained stable across the same age range (*p* = .986).

Taken together, these results demonstrate a clear cultural discrepancy in the developmental change of audiovisual integration. While Canadian infants exhibited a robust and stable capacity from 6 months onward, Chinese infants underwent a significant developmental progression, achieving a comparable level of performance only by 9 to 12 months of age. In addition to the interaction, the ANOVA also yielded a significant main effect of age group (*F*(1, 88) = 10.21, *p* = .002, *partial Eta Squared* = .11, *BF*_10_ = 13.31), which was primarily driven by the developmental change within the Chinese cohort. The main effect of country did not reach statistical significance (*F*(1, 88) = 0.97, *p* = .326, *partial Eta Squared* = .01, *BF*_10_ = 0.33).

The findings from the preceding experiment present an apparent paradox. While a cultural divergence in the onset of audiovisual integration was observed in early infancy, this difference appeared to resolve by 9 to 12 months of age, with both Canadian and Chinese infants demonstrating robust integration with familiar, same-race faces. This convergence in infancy seemingly contradicts a substantial body of literature on children and adults, which consistently documents persistent cross-cultural differences in audiovisual processing between Eastern and Western populations (Burnham & Lau, 1998; Sekiyama, 1997; Sekiyama & Burnham, 2008; Sekiyama & Tohkura, 1993).

This discrepancy raises a critical question: does the observed convergence reflect a genuine developmental parity, or does the relative perceptual ease of the task mask underlying cultural differences that manifest under more demanding conditions? To adjudicate between these possibilities, the subsequent experiment was designed to continue the exploration of cultural differences by testing the robustness of audiovisual integration. Specifically, we introduced a known perceptual challenge by using other-race faces as visual stimuli. This manipulation is based on prior work demonstrating that such unfamiliar stimuli can impair audiovisual integration, at least in East-Asian infants (Ujiie et al., 2020, 2021). By employing this more challenging paradigm, we aim to determine if the earlier maturation of this capacity in Western infants is also associated with greater resilience to perceptual interference, potentially revealing a more persistent cultural difference that reconciles our infant findings with the established literature.

## Experiment 3

### Participants

Twenty-one full-term English-learning White Canadian infants (10 females) with normal vision and hearing participated after caregivers provided informed consent. The participants were from 196 to 341 days old (*M* = 8.93 months, *SD* = 1.41 months). All participating infants were from monolingual families and had minimal exposure to East-Asian faces in daily life according to parental reports. To ensure that the other-race face manipulation in this experiment was valid (i.e., East Asian faces were indeed unfamiliar to the infants), we restricted recruitment to infants from households where both parents were White. Additional five infants’ data were excluded from data analyses due to fussiness (n = 2), parental interference (n = 2), the primary and/or secondary caregiver(s) was/were East-Asian (n = 1), and performance that was more than two standard deviations from the mean (n = 1). All participating families were from urban and suburban regions.

Thirty-three full-term Mandarin-learning Chinese infants (18 females) with normal vision and hearing participated after caregivers offered informed consent. The participants were from 183 to 380 days (*M* = 9.50 months, *SD* = 1.50 months). For this cohort, the concern regarding face-race exposure was naturally addressed by the region’s demographic composition: infants had negligible visual exposure to faces other than East Asian ones in daily life. Therefore, no infants were excluded based on caregivers’ race. Additional four infants’ data were removed because of calibration failure.

### Stimuli & Procedure

The experimental stimuli and procedures were consistent with those used in Experiment 2 except that White Canadian infants watched East-Asian female faces whereas Chinese participants watched White female faces.

## Results and Discussion

### Audiovisual integration

White Canadian infants completed 14.5 trials (*SD* = 2.48), while Chinese infants completed 8.45 trials (*SD* = 1.03). We conducted paired-sample *t*-tests on the participants average looking time in both types of trials. As shown in Figure 4a, the results indicated a significant preference for looking in Canadian infants (*t*(20) = 3.29, *p* = .004; Cohen’s *d* = 0.72, *BF*_10_ = 11.81), indicating that their auditory perceptual outcomes were significantly different between the two types of trials. In contrast to the findings in Experiment 2, Canadian infants exhibited a longer looking duration in the non-McGurk condition (*M* = 11.78, *SD* = 3.19) compared to the McGurk condition (*M* = 10.79, *SD* = 3.11) when presented with other-race faces. In contrast, Chinese infants showed comparable looking times in both types of trials (*M*_McGurk_ = 15.62, *SD*_McGurk_ = 2.65, *M*_non-McGurk_ = 15.10, *SD*_non-McGurk_ = 2.90; *t*(32) = 0.97, *p* = .340; Cohen’s *d* = 0.19, *BF*_10_ = 0.29), implying that they may not effectively integrate audiovisual speech syllables.

**Figure 4.**
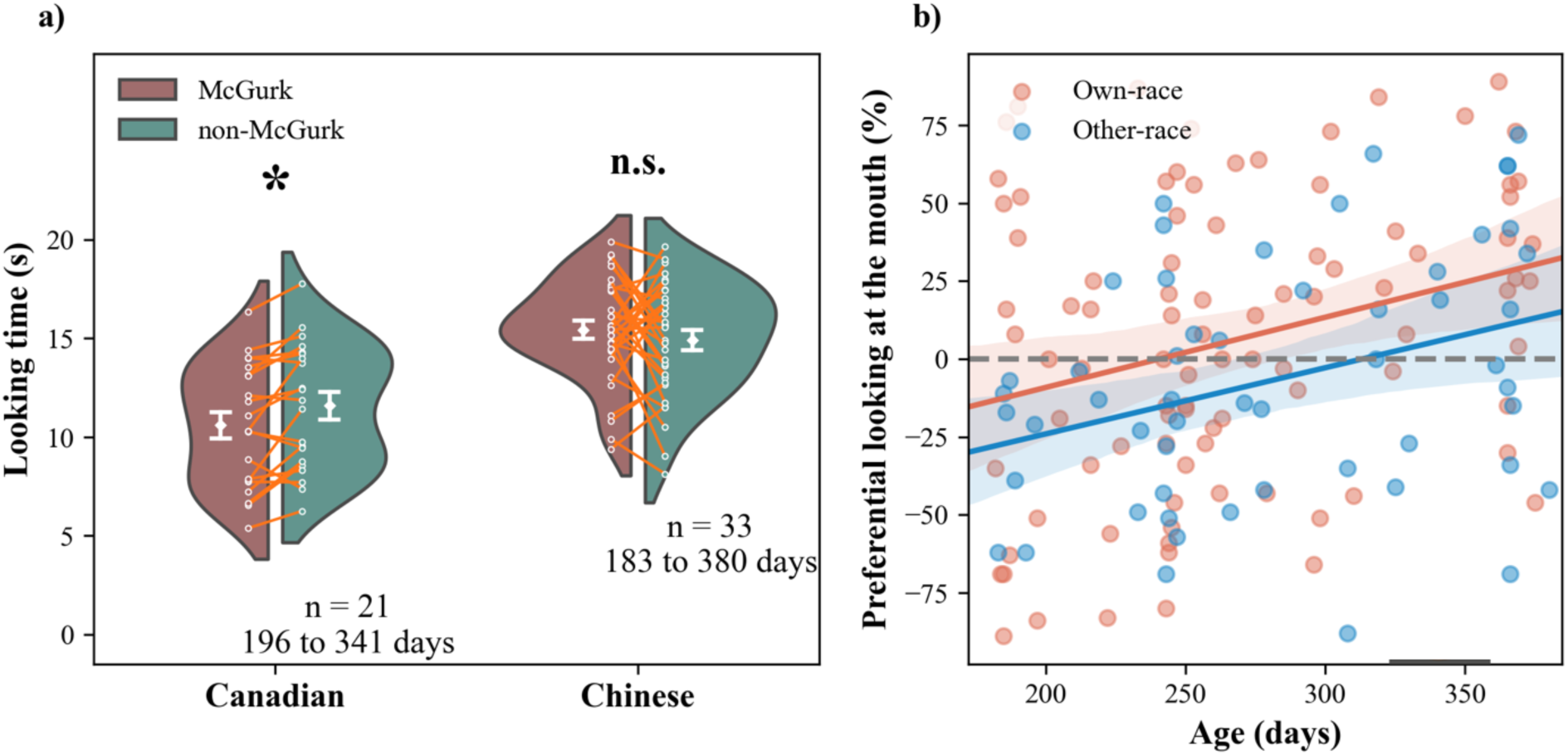
a) Mean looking time for the McGurk and non-McGurk trials in Canadian and Chinese infant cohorts in Experiment 3, in which infants watched other-race faces. The white diamond shapes represent the mean looking time. Error bars represent one standard error from the mean. Each participant’s looking time was annotated with tiny white dots, and the thin orange lines connecting the dots indicated the looking time of the same participant. The kernel distributions (the split violent shapes) represent the distribution of the mean looking time across participants. The asterisks indicate a statistically significant difference in looking time between the McGurk and non-McGurk trials. b) Eye-movement data from 2 experiments. Age-related change in the preferential looking time to the eye region (Eyes – Mouth) for own– and other-race faces. Data were collapsed from Asian and Western participants. Individual dots represent data from each participant. The shade areas around the trending lines represent one standard error. The horizontal dashed line indicates an equal looking time to the eyes and mouth regions. The AOIs for eyes and mouth regions were annotated as rectangle areas in the face image.

### Cultural difference in audiovisual integration

We used an independent-sample *t*-test to directly examine the cultural difference in audiovisual integration between the Asian and Western cohorts. Like the analysis performed in Experiment 2, we focused on the difference in looking time between the McGurk and non– McGurk trials as the dependent variable. The results showed a significant effect (*t*(52) = 2.11, *p* = .040; Cohen’s *d* = 0.59, *BF*_10_ = 1.67). This result suggested that even facing distraction from the presentation of other-race faces, Canadian infants still demonstrated stronger audiovisual integration than Chinese infants.

### Absence of age-related changes in both cohorts’ looking behaviors

We conducted Pearson’s correlation tests on infants’ age and looking time difference between two types of trials to confirm whether there were any developmental shifts underlying their looking patterns. We found no age-related change in looking behaviors in either cohort of participants (Canadian: *r*(19) = –.14, *p* = .555, *BF*_10_ = 0.54; Chinese: *r*(31) = .18, *p* = .308, *BF*_10_ = 0.60). These results indicated that when seeing other-race faces, Canadian infants looked consistently longer in the non-McGurk trials from 6 to 11 months of age, whereas Chinese infants remained not showing show any evidence of audiovisual integration across the 2nd half of infancy.

Together, these findings indicate that audiovisual integration, which was evident for own– race faces, was disrupted when infants viewed other-race faces. To examine whether this disruption was related to differences in visual attention to the face, we next analyzed infants’ face-scanning patterns. Specifically, we tested whether infants differed in the proportion of time spent looking at the mouth region, as attention to the mouth has been linked to enhanced audiovisual integration (e.g., Gurler et al., 2015; Stacey et al., 2020).

### Infants looked less at the mouth region of other-race faces across Experiments 2 and 3

To investigate the potential influence of face-scanning patterns on audiovisual integration, we analyzed infants’ preferential looking time to the mouth region relative to the eye region, a robust measure linked to speech and language development (Lewkowicz & Hansen– Tift, 2012). Areas of Interest (AOIs) for the eyes and mouth were defined using a face template approach, which ensures measurement comparability across different face stimuli by normalizing for variations in size, shape, or feature arrangement (Xiao & Lee, 2018; Xiao et al., 2026). For each infant, we calculated the proportional looking time to the mouth and eye AOIs relative to their total looking time to the face. The dependent variable for our analysis was the difference between these proportions (proportional mouth looking minus proportional eye looking).

We included data from Canadian and Chinese infants in Experiments 2 and 3, coding face race as own-race in Experiment 2 and other-race in Experiment 3. A linear regression model (see the formula below) was used to evaluate the effect of face race, country, age, and the interaction between face-race and country. As shown in Figure 4b, the results revealed a significant effect of age (*t* = 3.65, *p* < .001, *BF*_10_ = 79.95), infants increased their looking to the mouth region over the course of the first year of life. Moreover, we found a significant effect of face-race (*t* = 2.07, *p* = .040, *BF*_10_ = 1.34) infants looked less at the mouth region of other-race faces as compared to that of own-race faces. These results supported our prediction that infants, regardless of their cultural background, demonstrated distinctive looking patterns at other-race faces. This looking difference may further disrupt their ability to integrate audiovisual information effectively.

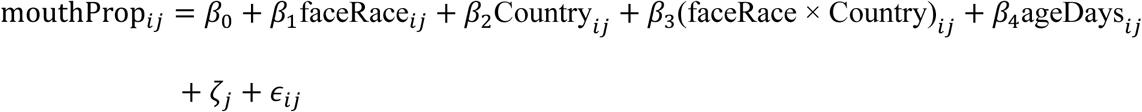

Crucially, the model revealed no significant main effect of cultural background, nor an interaction between cultural background and face race (all *t*s < 0.65, *p*s > .515, *BF*_10_s <= 0.36). This indicates that the face-scanning patterns, including the reduction in mouth-looking for other-race faces, were comparable across both Canadian and Chinese infants. Therefore, while the overall reduction in mouth-looking for other-race faces may contribute to the general difficulty of the task, this attentional factor does not appear to explain the specific cultural differences observed in audiovisual integration performance.

We also tested whether condition-specific mouth-looking predicted the strength of the McGurk effect. For each infant, we computed the difference in preferential mouth-looking between McGurk and non-McGurk trials and correlated this value with the corresponding McGurk-minus-non-McGurk looking-time difference. This analysis revealed no significant association, *r* = .03, *p* = .686, indicating that condition-related differences in mouth-looking did not predict the magnitude of audiovisual integration.

## General Discussion

Across three experiments, we found that audiovisual integration follows different developmental trajectories across cultures. White Canadian infants demonstrated robust integration by 6 months, whereas Chinese infants showed a more gradual emergence of integration across the first year. Although older Chinese infants reached comparable levels of integration with own-race faces, Experiment 3 revealed that this convergence was conditional: other-race faces disrupted integration in Chinese but not White Canadian infants. Thus, cultural differences persist not as a simple delay, but as differences in the robustness and generalizability of audiovisual integration.

We argue that these divergent timelines do not reflect developmental delay, but adaptive calibration to distinct communicative ecologies (Bornstein, 2013; Werker & Hensch, 2015). Here, “input” refers broadly to infants’ everyday speech and socio-communicative environment, including the diversity of talkers, phonological and prosodic variability, and caregiver–infant interaction patterns. White Canadian infants may specialize earlier under relatively consistent input, whereas Chinese infants may maintain greater flexibility while adapting to more variable communicative environments.

### The roots of early divergence: caregiving and linguistic environments

What drives this initial 6-month divergence? We propose two macro-level environmental factors: caregiver consistency and linguistic structure. First, the two cohorts experience fundamentally different social networks. Our demographic data show that the Canadian infants were raised exclusively by their mothers (100% sole primary caregiver). The Chinese infants experienced distributed caregiving, with extended family (e.g., grandmothers) acting as primary caregivers in 39% of families. The number of talkers in an infant’s environment directly structures their moment-to-moment language exposure (Bunce et al., 2025). By interacting with a larger, more variable network of communication partners, Chinese infants undergo early “perceptual variability training.” This forces their developing system to remain flexible longer, delaying the onset of rigid audiovisual binding.

Our demographic data suggest that the two cohorts differed in early caregiving environments. White Canadian infants were primarily cared for by mothers, whereas Chinese infants more often experienced distributed caregiving involving extended family members. Such differences may alter the diversity and variability of infants’ everyday communicative input, potentially shaping the timing with which audiovisual correspondences are learned.

Second, the structural properties of the ambient languages differ. English is stress-timed, featuring rhythmic variability, frequent vowel reduction, and complex consonant clusters. These acoustic properties make visual speech cues highly informative for parsing the speech stream (Birules et al., 2018), potentially driving English-learning infants to rely earlier and more heavily on lip movements. Mandarin is syllable-timed, with uniform rhythm and a fundamental reliance on pitch contours (lexical tone) that lack clear visual correlates.

However, comparing our findings with the broader literature introduces an important caveat: Japanese infants and children also exhibit delayed audiovisual integration compared to Western cohorts (e.g., Geangu et al., 2016; Haensel et al., 2020; Sekiyama & Burnham, 2008), despite Japanese being a mora-timed language that is rhythmically distinct from both English and Mandarin. This convergence between Mandarin and Japanese learners suggests that language rhythm alone cannot explain the East-West divide. Shared sociocultural practices likely play an independent, compounding role. Because both Mandarin and Japanese lack the consonant clusters of English, a linguistic contribution cannot be entirely ruled out, but it must interact with these broader socio-communicative norms.

### Mechanisms of convergence: the 6-to-12-month shift

If Chinese infants navigate a more variable social network and face different linguistic demands, what experiential shifts enable them to catch up by 12 months? We propose a cascade of three interacting developmental processes: cumulative statistical learning, stabilization of native phonology, and the onset of motor practice.

First, differences in maternal input may demand different accumulation thresholds. Western caregivers typically use infant-directed speech (IDS) heavily marked by hyper– articulated visual cues (Shochi et al., 2009). This high-contrast signal acts as a potent tutor, accelerating cross-modal binding. If East Asian communicative norms favor less exaggerated visual prosody, the signal-to-noise ratio is lower. Chinese infants may simply require a larger cumulative corpus of input, gathered over the second half of the first year, to extract reliable viseme-phoneme statistics from their diverse caregivers.

Second, this period coincides with perceptual narrowing. For infants acquiring a tonal language, a critical task between 6 and 9 months is stabilizing lexical pitch categories (Mattock & Burnham, 2006). Because pitch lacks a salient visual correlate on the lips, Mandarin learners must prioritize auditory tracking over visual speech cues. Once these tonal categories stabilize late in the first year, cognitive and attentional resources are freed to process the segmental visual cues necessary for the McGurk effect.

Third, this timeline overlaps perfectly with the onset of canonical babbling. Active articulation allows infants to build internal sensorimotor forward models, physically mapping specific mouth movements to their acoustic consequences (Bruderer et al., 2015). For Chinese infants relying on variable external input, their own self-generated motor practice may serve as an internal catalyst that finally prepares audiovisual integration by 12 months.

### Conditional Convergence and Its Implications

Our findings reveal a conditional, rather than complete, convergence in audiovisual integration. As shown in Experiment 3, when other-race faces were introduced, cultural differences re-emerged. White Canadian infants continued to show reliable integration, whereas Chinese infants, including those in the older age range, no longer exhibited a reliable McGurk effect. Thus, cultural differences do not simply disappear by 12 months; rather, audiovisual integration becomes broadly available under familiar conditions but differs in robustness and generalizability when perceptual demands increase.

This conditional pattern helps reconcile the infant findings with evidence from older children and adults. Adult studies typically show that individuals across cultures can experience the McGurk effect, but differ in the magnitude of susceptibility or in the relative weighting of auditory and visual speech cues. Our findings suggest that these later cultural differences may reflect long-term consequences of early differences in how audiovisual integration generalizes across perceptual contexts, rather than contradicting the apparent convergence observed under familiar own-race conditions in infancy.

### Attentional mechanisms and stimulus constraints

Prior research suggests that Western infants attend to the mouth earlier and longer than East Asian infants (Geangu et al., 2016; Haensel et al., 2020), making visual attention a potential mechanism underlying early cultural divergence. However, our eye-tracking data did not support this account: we found no cultural differences in mouth-looking, and mouth-looking did not predict the strength of the McGurk effect.

This null result may reflect properties of our stimulus design. Because faces alternated between articulating and static states every second, the onset of lip movement on an otherwise stationary face may have strongly attracted bottom-up attention to the mouth across infants, reducing variability in spontaneous face-scanning strategies. Experiment 3 provides further support for this interpretation: when infants viewed other-race faces, mouth-looking decreased across both cohorts and the McGurk effect was reduced correspondingly. Thus, mouth-directed attention appears important for audiovisual integration, but our paradigm may have limited sensitivity to cultural differences in naturalistic scanning patterns.

Future studies using continuous and more naturalistic speech stimuli will be needed to determine whether culturally shaped differences in spontaneous face scanning contribute to the developmental trajectory of audiovisual integration.

### Paradigm sensitivity and stimulus validity

The cultural divergence observed here cannot be explained by testing Chinese infants with unfamiliar speech sounds. Our stimuli used basic plosive consonants (/b/, /p/, /g/, and /k/) paired with /a/, which are common contrasts in both English and Mandarin. Moreover, a recent multinational study showed comparable McGurk susceptibility for these contrasts among Mandarin– and English-speaking adults (Magnotti et al., 2025). Thus, the developmental difference observed here likely reflects variation in the emergence of cross-modal integration rather than acoustic unfamiliarity.

Our findings also extend previous work suggesting that cultural differences in the McGurk effect emerge only later in childhood (Sekiyama & Burnham, 2008). Unlike explicit categorization tasks, which require participants to assign a phonological label to an illusory percept, our looking-time paradigm provides an implicit measure of audiovisual binding. By assessing whether visual speech information alters infants’ perception of auditory sequences, this approach reveals that experience-related differences in multisensory integration can emerge within the first year of life.

Finally, our crossover design reduces the possibility that the observed effects reflect low– level preferences for particular sounds or faces. Because the same audiovisual tokens were presented across conditions, differences in looking behavior depended on the relationship between auditory and visual information rather than stimulus salience alone. Experiment 3 further demonstrated that identical audiovisual inputs produced different integration patterns depending on face-race context, supporting the role of experience-dependent perceptual processes.

### Study limitations and future directions

We note several limitations. First, our cross-sectional design captures group-level patterns; longitudinal tracking is required to map individual developmental trajectories and establish causal links between specific environmental inputs and integration outcomes. Second, while we utilized shared phonemic contrasts to ensure valid comparison, future designs should probe language-specific McGurk stimuli (e.g., using contrasts unique to Mandarin) to test how fine-grained ambient statistics modulate integration. Third, although parents were instructed to remain neutral, they were not blinded to the stimuli, leaving room for unconscious cueing. Future protocols should mandate opaque partitions and sound-masking headphones for caregivers. Finally, our data reflect urban, educated populations in Canada and China. To uncouple the intersecting roles of language rhythm and social environment, future research must sample more broadly by testing infants acquiring syllable-timed Western languages (e.g., Spanish or French) and stress-timed Asian languages.

