## supplementalMaterials for "Early Emergence of Cultural Differences in Audiovisual Speech Perception"

**Supplemental Materials: Caregiving arrangement**

**Methods**

Data on caregiving arrangements were collected via a parental questionnaire administered to the families of all participating infants in both Canada and China. Caregivers were asked to detail the infant's daily face-to-face interactions. The predefined list of interaction partners included: Mother, Father, Grandmother, Grandfather, Nanny/Babysitter, Older Brother, Older Sister, and an open field for "Other" regular contacts.

For each individual interaction partner, caregivers were required to report the following specific metrics:

- **Language Spoken:** Coded as 1 (Native language: Mandarin or English, depending on the cohort) or 2 (Other language).
- **Race / Ethnicity:** The racial or ethnic background of the interaction partner.
- **Age:** The age of the interaction partner.
- **Contact Duration:** The estimated daily duration of interaction involving direct face-to-face contact with the infant.
- **Direct Speech Duration:** The estimated daily duration the partner spent speaking directly to the infant.

**Results**

The analysis of questionnaire responses revealed significant cross-cultural variations in caregiving practices between the Canadian and Chinese cohorts across three primary domains.

***Parents and Grandparents Involvement***

A significant difference was observed in the number of hours parents were directly involved in caregiving. In the Canadian cohort, mothers and fathers reported a combined average of 29.6 hours (*SD* = 11.38) of caregiving per day. In contrast, Chinese parents reported a significantly lower combined average of 9.5 hours (*SD* = 4.69) per day. This difference was largely accounted for by greater involvement from alloparents, particularly grandparents, in the Chinese cohort. Chinese grandparents were reported to provide an average of 4.8 hours (*SD* = 5.04) of daily care, more than double the 2.0 hours (*SD* = 0.86) reported for Canadian grandparents.

***Primary Caregiver Identity***

The identity of the primary caregiver differed starkly between the two cultures. In the Canadian sample, mothers were exclusively identified as the primary caregiver in 100% of participating families. In the Chinese sample, caregiving roles were more distributed. Mothers were identified as the primary caregiver in only 43.7% of families, while grandmothers assumed this role nearly as often (39.1%). Other primary caregivers in China included nannies (2.3%) and grandfathers (1.5%).

***Shared Caregiving Responsibilities***

In the Canadian cohort, caregiving duties were consistently reported as being shared between the mother and father. The Chinese cohort, however, showed a more varied distribution of shared responsibilities that reflected the distributed caregiving model. While some families reported shared care between the father and mother (5.7%), other common arrangements included sharing between the mother and grandmother (4.6%), the father and grandfather (2.3%), or the mother and a nanny (1.5%).

These findings provide quantitative evidence of two distinct caregiving models within our samples: a nuclear-family-centric model in the Canadian cohort and a distributed, multi-generational model in the Chinese cohort.

**Supplemental Material: Validation of McGurk Stimuli**

To verify that the specific McGurk stimuli and the unique alternating “moving-still” presentation pattern utilized in the infant experiments reliably elicit audiovisual speech integration, we conducted a behavioral validation study with adult participants.

**Participants**

A total of 32 Chinese university students (13 males; mean age = 18.9 years, *SD* = 0.86, range: 18–21 years) participated in this study. All participants were right-handed, possessed normal or corrected-to-normal visual and auditory acuity, participated voluntarily, and provided written informed consent prior to the experiment. Each participant received monetary compensation upon completion of the study.

**Stimuli and Apparatus**

To ensure that the adult validation accurately mirrored the perceptual experience of the infant paradigm, the visual stimuli were face videos used in the main experiments. Audiovisual stimuli were generated by pairing silent articulatory video clips with auditory syllables.

The stimulus set included **congruent pairings** (matching audio and visual syllables, e.g., /ba/ visual–/ba/ audio), **McGurk incongruent pairings** designed to elicit audiovisual integration (e.g., /ga/ visual–/ba/ audio; /ka/ visual–/pa/ audio), and **non-McGurk incongruent pairings** designed to create audiovisual conflict without fusion (e.g., /ba/ visual–/ga/ audio; /pa/ visual–/ka/ audio). These pairings were generated across two face races (East Asian and White). All stimuli were presented on a 17-inch monitor, and audio output was calibrated to a comfortable listening level of 75–80 dB.

**Design and Procedure**

The experiment employed a 2 (Face Race: East Asian vs. White) × 2 (Interference Type: Audiovisual Integration vs. Audiovisual Conflict) within-subjects design. The dependent measure was auditory response accuracy, which served as an index of how strongly the incongruent visual information interfered with veridical auditory speech perception. The experiment was programmed and executed using PsychoPy (Version 2024.1.4) in a quiet laboratory setting.

Participants completed two blocks of a forced-choice identification task, corresponding to the two syllable sets (/ka/-/pa/ and /ga/-/ba/). Each trial began with a 500 ms central fixation cross, followed by a 1 s audiovisual syllable stimulus. Participants were instructed to attend carefully to both the visual and auditory components. Following stimulus offset, a response screen prompted participants to indicate the consonant they perceived via a 6-alternative forced-choice keypress (“g”, “b”, “d”, “k”, “p”, or “t”), followed by a 5-point confidence rating.

Block order was counterbalanced across participants. Within each block, face race, audiovisual congruency, and sound type were fully randomized. Each block contained 64 trials, yielding a total of 128 trials per participant. Prior to the main experiment, participants completed 4 practice trials to familiarize themselves with the task demands and response mappings.

**Data Analysis**

The primary dependent variable was the *interference effect*, defined as the reduction in auditory identification accuracy caused by incongruent visual information relative to a congruent baseline. To calculate this, baseline accuracy was first established using the congruent trials. The **audiovisual conflict effect** (interference elicited by non-McGurk stimuli) was calculated as the accuracy for congruent trials minus the accuracy for non-McGurk incongruent trials. The **audiovisual integration effect** (interference elicited by McGurk stimuli) was calculated as the accuracy for congruent trials minus the accuracy for McGurk incongruent trials.

To focus on the core theoretical factors and simplify the analysis, the interference scores were collapsed across the two syllable pairings (/ka/-/pa/ and /ga/-/ba/). The data were then submitted to a 2 (Face Race: East Asian, White) × 2 (Interference Type: Audiovisual Conflict, Audiovisual Integration) repeated-measures ANOVA.

**Results**

To confirm the McGurk illusion, we examined the magnitude of visual interference on auditory perception. The repeated-measures ANOVA revealed a significant main effect of Interference Type, *F*(1, 31) = 16.83, *p* < .001. As hypothesized, the McGurk stimuli (Audiovisual Integration) produced a substantially larger overall interference effect (*M* = 29.3%) than the non-McGurk control stimuli (Audiovisual Conflict, *M* = 13.7%). The main effect of Face Race was not significant, *F*(1, 31) = 2.90, *p* = .098.

Moreover, the analysis revealed a significant interaction between Face Race and Interference Type, *F*(1, 31) = 4.55, *p* = .041. To decompose this interaction, Bonferroni-adjusted simple main effect analyses were conducted to compare the magnitude of Integration versus Conflict interference within each face race. For East Asian faces, the audiovisual integration effect (*M* = 32.3%, *SE* = 5.3%) was significantly larger than the audiovisual conflict effect (*M* = 13.4%, *SE* = 4.3%), *t*(31) = 4.43, *p* < .001. For White faces, the audiovisual integration effect (*M* = 26.2%, *SE* = 5.1%) was also significantly larger than the audiovisual conflict effect (*M* = 13.9%, *SE* = 4.3%), *t*(31) = 3.12, *p* = .004.

This adult validation experiment confirmed that the specific synthetic stimuli used in the infant studies reliably elicit robust audiovisual speech integration. The significant interaction indicates that the magnitude of the McGurk illusion was slightly more pronounced when Chinese adults viewed own-race (East Asian) faces compared to other-race (White) faces. These findings definitively validate the effectiveness of the stimulus design and support its application in measuring cross-modal speech integration in our infant cohorts.

**Supplemental Material: Validation of Synthetic Talking Faces**

To assess the perceived naturalness of the GAN-generated synthetic talking faces used in the main study, we recruited a separate cohort of sixteen Asian undergraduate students (12 female; mean age = 19 years) from a Chinese university. All participants reported having normal or corrected-to-normal vision and had no prior exposure to the talking face stimuli used in the main study. The study protocol was approved by the local Human Research Ethics Committee, and written informed consent was obtained from all participants prior to data collection. The stimuli set comprised 80 dynamic talking-face videos, equally distributed between Real talking faces recorded from human individuals and Synthetic talking faces generated using DeepFaceLab. Although the main study utilized a broader set of 16 actresses (8 Asian and 8 Caucasian), the current validation relied on a subset of 10 female actresses (5 Asian and 5 Caucasian). This selection was restricted to these specific actresses because they possessed corresponding real-world video recordings of infant-directed speech, which were required to serve as the ground-truth baseline for the "Real versus Synthetic" comparison. All videos were presented in color against a gray background with consistent frontal-view framing. To present the diverse profile of each actress, we extracted 4 3-seconds clips from the video, leading to 4 clips $\times$ 2 faces-races $\times$ 5 actresses $\times$ 2 formats $=$ 80 video stimuli.

Following a practice block to familiarize participants with the task requirements, the main experiment consisted of 80 trials presented in a randomized order. Each trial began with a central fixation cross displayed for 500 ms, followed by the presentation of a talking-face video for a fixed duration of 3000 ms. Immediately upon the offset of the video, participants were required to perform two consecutive tasks in a fixed order. First, they completed a race classification task, a two-alternative forced-choice (2AFC) paradigm, in which they identified the face as either “Asian” or “White”. This was followed by a naturalness rating task, where participants evaluated the authenticity of the face on a 7-point Likert scale ranging from 1 (“Very Unnatural”) to 7 (“Very Natural”). This dual-task design ensured that participants were actively attending to the visual-structural information of the faces before providing subjective ratings.

**Results**

The primary objective of this validation was to determine if the synthetic videos were perceived as naturally as the real footage. We first analyzed the naturalness ratings to establish a baseline of validity. A one-sample *t*-test against the scale midpoint of 4 (representing "neutral/uncertain") demonstrated that naturalness ratings were significantly above the midpoint for both real faces (*M* = 4.89, *SD* = 0.92, *p* = .002, Cohen's *d* = 0.96) and synthetic faces (*M* = 4.71, *SD* = 0.97, *p* = .011, Cohen's *d* = 0.73), indicating that the generated stimuli were explicitly perceived as natural rather than artificial. Crucially, a direct comparison between the two conditions revealed no statistically significant difference in naturalness ratings between real and synthetic faces (*t*(15) = 1.42, *p* = .18, Cohen's *d* = 0.36). Bayesian analysis yielded a Bayes Factor (*BF*_01_) of 1.68, providing positive support for the null hypothesis. These findings suggest that the synthetic faces were perceptually indistinguishable from the real human recordings, thereby meeting the standard for valid experimental stimuli.

Given the cross-cultural context of the main study, it was also essential to verify that the quality of face generation was consistent across racial groups. We compared the naturalness ratings for the generated faces and found no significant difference between synthetic Asian and synthetic White faces (*t*(15) = 1.12, *p* = .28, Cohen's *d* = 0.28), confirming their validity for use in the main experiments.

Finally, we examined the performance on the race classification task to verify participant engagement and the preservation of racial features in the synthetic stimuli. Participants exhibited near-ceiling accuracy for both real faces (*M* = 0.97, *SD* = 0.03) and synthetic faces (*M* = 0.95, *SD* = 0.06). A paired-sample *t*-test confirmed there was no significant difference in classification accuracy between the face types (*t*(15) = 1.41, *p* = .18, Cohen's *d* = 0.35). This high level of accuracy provides robust evidence that the participants were attentive to the task and that the synthetic generation process successfully retained distinct, identifiable racial information even when only face-internal information was presented.

**Supplemental Analysis: Trial-Level Dynamics and Exposure Controls for Experiment 2**

Experiment 2 showed differences between White Canadian and Chinese infants in both the number of usable trials and the average looking time per trial. These differences raise an important methodological question: could variation in trial completion, average looking duration, habituation, or accumulated exposure account for the observed Country × Age Group × Trial Type pattern in audiovisual integration?

We addressed this question using three complementary trial-level linear mixed-effects models and an exposure-matched sensitivity analysis. Linear mixed-effects models were fitted in R (Version 4.6.1) using lme4 (Version 2.0.1) and lmerTest (Version 3.2.1). Type III tests were obtained using Satterthwaite’s approximation after setting sum-to-zero contrasts (options(contrasts = c("contr.sum", "contr.poly"))). The first model tested whether the key Country × Age Group × Trial Type effect remained when trial progression was included as a covariate. The second model tested whether the rate of within-session decline in looking time differed between cultural cohorts. The third model tested whether the McGurk versus non-McGurk difference changed as a function of trial progression. Finally, the exposure-matched analysis tested whether the main developmental pattern remained when cumulative looking time was equated across groups.

**Model 1: Primary Trial-Level Control Model**

The first model tested whether the central Country × Age Group × Trial Type effect remained after accounting for trial progression. Total looking time on each trial was predicted from Country, Age Group, Trial Type, and log-transformed trial number (logTrial). The model included all interactions among Country, Age Group, and Trial Type, with participant identity included as a random intercept.

The model specification was:

totalLooking ~ Country * AgeGroup * TrialType + logTrial + (1 | ParticipantID)

where totalLooking refers to looking duration on each trial, Country refers to the cultural cohort (White Canadian vs. Chinese), AgeGroup refers to younger versus older infants, TrialType refers to McGurk versus non-McGurk trials, and logTrial captures trial progression across the experimental session.

A Type III ANOVA using Satterthwaite’s method produced the results shown in Table S1.

**Table S1**
*Trial-level mixed-effects model testing the Country × Age Group × Trial Type effect while controlling for trial progression.*

| Effect | F | *p* |
| --- | --- | --- |
| Country | 3.9 | 0.051 |
| Age Group | 0.22 | 0.638 |
| Trial Type | 0.91 | 0.341 |
| logTrial | 328.13 | < .001 |
| Country × Age Group | 0.26 | 0.611 |
| Country × Trial Type | 0.4 | 0.526 |
| Age Group × Trial Type | 14.13 | < .001 |
| Country × Age Group × Trial Type | 11.05 | < .001 |

The main effect of logTrial was significant, indicating that looking time declined over the course of the experiment, consistent with general habituation or fatigue. Critically, the Country × Age Group × Trial Type interaction remained significant after controlling for trial progression. This indicates that the key cultural-developmental pattern was not reducible to overall changes in looking time across the session.

The marginal main effect of Country should not be interpreted in isolation because the significant three-way interaction indicates that cultural differences in looking behavior depended jointly on age group and trial type.

**Model 2: Cross-Cultural Trial-Progression Slope Check**

The second model tested whether the rate of within-session decline differed between White Canadian and Chinese infants. This model added the Country × logTrial interaction to the primary model. This analysis directly addresses whether the two cultural cohorts differed in habituation or fatigue slopes across trials.

The model specification was:

totalLooking ~ Country * AgeGroup * TrialType + Country:logTrial + logTrial + (1 | ParticipantID)

A Type III ANOVA using Satterthwaite’s method produced the results shown in Table S2.

**Table S2**
*Trial-level mixed-effects model testing whether trial-progression slopes differed by cultural cohort.*

| Effect | F | *p* |
| --- | --- | --- |
| Country | 2.55 | 0.111 |
| Age Group | 0.22 | 0.636 |
| Trial Type | 0.91 | 0.34 |
| logTrial | 308.16 | < .001 |
| Country × Age Group | 0.26 | 0.613 |
| Country × Trial Type | 0.41 | 0.524 |
| Age Group × Trial Type | 14.13 | < .001 |
| Country × logTrial | 0.02 | 0.883 |
| Country × Age Group × Trial Type | 11.06 | < .001 |

The main effect of logTrial was again significant, confirming an overall decline in looking time across trials. However, the Country × logTrial interaction was not significant, indicating that the rate of within-session decline did not differ between White Canadian and Chinese infants. Thus, the difference in completed trial numbers cannot be attributed to different habituation or fatigue slopes across the two cultural cohorts.

Importantly, the Country × Age Group × Trial Type interaction remained significant in this slope-check model, confirming that the main cultural-developmental pattern was preserved even when cultural differences in trial-progression slopes were explicitly tested.

**Model 3: Trial Type × Trial Progression Check**

Anonymous reviewer raised the possibility that infants, particularly Chinese infants, might require more accumulated exposure within the experimental session before exhibiting a McGurk effect. A simple first-half versus second-half division is difficult to define consistently because infants contributed different numbers of valid trials depending on individual habituation and session completion. Such a division is also difficult to interpret because trial order varied in syllables and face identities. We therefore tested this question using trial progression as a continuous predictor.

This model tested whether the McGurk versus non-McGurk difference changed across the experimental session by adding the Trial Type × logTrial interaction to the primary trial-level model.

The model specification was:

totalLooking ~ Country * AgeGroup * TrialType + TrialType:logTrial + logTrial + (1 | ParticipantID)

A Type III ANOVA using Satterthwaite’s method produced the results shown in Table S3.

**Table S3**
*Trial-level mixed-effects model testing whether the McGurk versus non-McGurk difference changed as a function of trial progression.*

| Effect | F | *p* |
| --- | --- | --- |
| Country | 3.83 | 0.053 |
| Age Group | 0.24 | 0.625 |
| Trial Type | 0.29 | 0.59 |
| logTrial | 329.18 | < .001 |
| Country × Age Group | 0.26 | 0.611 |
| Country × Trial Type | 0.15 | 0.702 |
| Age Group × Trial Type | 14.31 | < .001 |
| Trial Type × logTrial | 1.03 | 0.31 |
| Country × Age Group × Trial Type | 11.2 | < .001 |

The Trial Type × logTrial interaction was not significant, indicating that the McGurk versus non-McGurk difference did not systematically increase or decrease over the course of the experiment. Thus, there was no evidence that the McGurk effect emerged only after infants accumulated more exposure within the experimental session. At the same time, the Country × Age Group × Trial Type interaction remained significant, confirming that the main cultural-developmental pattern was preserved when trial-level changes in the McGurk effect were explicitly tested.

**Shorter Per-Trial Looking Did Not Prevent Detection of Condition Effects**

One concern is that shorter looking times per trial might reduce sensitivity to detect McGurk versus non-McGurk differences. This prediction was not supported by the observed data. White Canadian infants showed shorter average looking times per trial than Chinese infants, yet White Canadian infants exhibited a significant McGurk versus non-McGurk effect. Thus, shorter per-trial looking time did not prevent detection of condition differences in the group for which this concern would be most relevant.

**Exposure-Matched Sensitivity Analysis**

To further evaluate whether differences in accumulated exposure could account for the findings, we conducted an exposure-matched sensitivity analysis. Because White Canadian infants completed more trials on average but looked for less time within individual trials, we truncated the White Canadian dataset to the first 12 trials. This produced total accumulated looking time comparable to the full Chinese dataset.

The exposure-matched comparison yielded statistically equivalent cumulative exposure between the White Canadian subset and the Chinese cohort (White Canadian 12-trial subset: $M=147.3$ s; Chinese full dataset: $M=140.6$ s), $t(89.43)=0.66,p=.51$.

Within this exposure-matched White Canadian subset, the McGurk versus non-McGurk effect remained significant, $t(44)=2.95,p=.005$.

We then repeated the 2 (Country) × 2 (Age Group) ANOVA on McGurk-minus-non-McGurk difference scores using the exposure-matched White Canadian data. The critical interaction remained significant ($F(1,88)=11.72,p<.001$), as did the Age Group main effect ($F(1,88)=10.17,p=.002$), whereas the Country main effect was not significant ($p=.36$).

These results indicate that the main cultural-developmental pattern remained after cumulative exposure was equated across groups.

**Relation to Benchmark Experiments**

The expanded benchmark data from Experiments 1a and 1b revealed a similar pattern of cross-cultural differences in overall looking behavior, despite the absence of audiovisual conflict or McGurk stimuli. Across both benchmark experiments, Chinese infants consistently exhibited longer average looking times per trial than White Canadian infants (Experiment 1a: 14,516 vs. 13,754 ms; Experiment 1b: 15,897 vs. 10,329 ms). In contrast, the average number of completed trials showed no consistent cultural pattern: the two cohorts completed comparable numbers of trials in Experiment 1a (White Canadian: 7.96; Chinese: 7.75), whereas White Canadian infants completed more trials than Chinese infants in Experiment 1b (13.48 vs. 10.40).

These findings indicate that within-trial looking duration and the number of completed trials capture distinct aspects of infants’ behavior and do not reflect a single dimension of engagement or task compliance. More importantly, because these benchmark experiments involved only auditory sequence processing and contained no McGurk manipulation, the observed cross-cultural differences in overall looking behavior cannot explain the condition-specific developmental pattern of audiovisual integration observed in Experiment 2.

**Conclusion**

Across analyses, differences in usable trial counts, average per-trial looking duration, trial progression, habituation slopes, within-session changes in the McGurk effect, and total accumulated exposure did not account for the observed cultural-developmental pattern in Experiment 2. The key Country × Age Group × Trial Type interaction remained significant after controlling for trial progression, after explicitly testing the Country × logTrial slope, and after testing whether the Trial Type × logTrial interaction captured within-session emergence of the McGurk effect. In addition, the Country × Age Group interaction remained significant in the exposure-matched sensitivity analysis. Therefore, the developmental divergence in audiovisual integration cannot be attributed to differences in trial architecture, looking-time duration, or accumulated exposure.
